# Predicting Inhibitors of OATP1B1 via Heterogeneous OATP-Ligand Interaction Graph Neural Network (HOLIgraph)

**DOI:** 10.1101/2024.10.01.615464

**Authors:** Mehrsa Mardikoraem, Joelle N. Eaves, Theodore Belecciu, Nathaniel Pascual, Alexander Aljets, Bruno Hagenbuch, Erik M. Shapiro, Benjamin J. Orlando, Daniel R. Woldring

**Author notes:** These authors contributed equally to this work (Mehrsa Mardikoraem, Joelle N. Eaves). Contributing authors.

## Abstract

Organic anion transporting polypeptides (OATPs) are membrane transporters crucial for drug uptake and distribution in the human body. OATPs can mediate drug-drug interactions (DDIs) in which the interaction of one drug with an OATP impairs the uptake of another drug, resulting in potentially fatal pharmacological effects. Predicting OATP-mediated DDIs is challenging, due to limited information on OATP inhibition mechanisms and inconsistent experimental OATP inhibition data across different studies. This study introduces Heterogeneous OATP-Ligand Interaction Graph Neural Network (HOLIgraph), a novel computational model that integrates molecular modeling with a graph neural network to enhance the prediction of drug-induced OATP inhibition. By combining ligand (i.e., drug) molecular features with protein-ligand interaction data from rigorous docking simulations, HOLIgraph outperforms traditional DDI prediction models which rely solely on ligand molecular features. HOLIgraph achieved a median balanced accuracy of over 90 percent when predicting inhibitors for OATP1B1, significantly outperforming purely ligand-based models. Beyond improving inhibition prediction, the data used to train HOLIgraph can enable the characterization of protein residues involved in inhibitory drug-OATP interactions. We identified certain OATP1B1 residues that preferentially interact with inhibitors, including I46 and K49. We anticipate such interaction information will be valuable to future structural and mechanistic investigations of OATP1B1.

**Scientific Contribution:** HOLIgraph introduces a new paradigm for DDI prediction by incorporating protein-ligand interactions derived from docking simulations into a graph neural net framework. This approach, enabled by recent structural breakthroughs for OATP1B1, represents a significant departure from traditional models that rely only on ligand features. By achieving high predictive accuracy and uncovering mechanistic insights, HOLIgraph sets a new trajectory for computational tools in drug design and DDI prediction.

## Introduction

Organic anion transporting polypeptides (OATPs) are a family of membrane proteins responsible for transporting a wide range of drugs and endogenous compounds into the liver and other pharmacologically relevant tissues (e.g., kidneys and intestines) [1]. To accommodate the variety of substrates processed by these tissues, OATPs exhibit a high degree of promiscuity [2]. This promiscuity gives rise to intricate intermolecular interactions and elaborate regulatory networks that can profoundly influence drug efficacy and toxicity. The US Food and Drug Administration (FDA) recognizes hepatic OATP1B1 and OATP1B3 as common participants in clinical drug-drug interactions. As such, FDA guidance suggests preclinical investigations to identify how a new drug candidate interacts with these hepatic OATPs [3].

Understanding the transport and inhibition mechanisms of OATPs and other membrane transport proteins is particularly challenging since they are inherently difficult to isolate for structural determination and functional characterization. In the past two decades, however, computational molecular modeling techniques have greatly expedited and improved functional characterization efforts like transporter inhibition prediction [4]. There are two types of computational molecular models typically implemented to predict whether a ligand (drug) will inhibit a transporter protein: ligand-based and structure-based models [2].

Ligand-based models relate the physical structure and chemical properties of a ligand to its empirical transporter activity. Ligand-based models are trained on experimental datasets consisting of large numbers of ligands, learning trends between physicochemical ligand properties and transporter activity. Trained models can then predict how a new ligand will interact with the transporter of interest through extrapolation of these learned trends. For OATP inhibition prediction, many ligand-based models have been developed with good predictive power [5–8]. However, ligand-based models generally provide limited mechanistic insight, and may only use a small fraction of the physicochemical information relevant for determining inhibition or transport [2]. It is well known that interactions between specific OATP residues and ligands play a role in ligand-transporter behavior [9], thus we hypothesized that leveraging such information in addition to traditional ligand-only information may enhance model prediction performance.

Structure-based models contain information about the bound protein-ligand complex, typically involving features like the binding interface energy or molecular docking simulation scores [10]. Recently, attention has been drawn toward interaction-based modeling, which considers more intricacies of the protein-ligand binding interface.

For example, the innovative incorporation of interaction-information into classification models by Aniceto et al. [10] has shown great promise in predicting urease inhibitors. Interaction data, such as the type and number of interactions in the protein-ligand binding interface, enabled a promising 80% precision in their urease classifier model and offered substantial insight into residue-importance in urease inhibition [10]. Information encoded in interaction-based models may be obtained from experimentally resolved protein structures, either experimentally elucidated in complex with the lig- and, or complexed via molecular docking simulations. It is also possible to obtain interaction data from computationally predicted structures (e.g., structures predicted by AlphaFold), though this is not recommended due to observed performance losses [11, 12]. The recent publication of multiple high-resolution cryo-EM structures of OATP1B1 (apo, holo, inward-facing, outward-facing) greatly facilitates the obtainment of high-quality interaction data between OATP1B1 and its ligand partners [9].

We present a novel approach to integrate both ligand- and structure-based modeling into a single holistic model that greatly expands upon the predictive power of conventional classifier models: HOLIgraph (Heterogeneous OATP-Ligand Interaction Graph Neural Network). HOLIgraph employs state-of-the-art graph neural network (GNN) technology to correlate ligand activity to ligand physicochemical and topological properties, protein sequence, and detailed protein-ligand interaction information obtained via rigorous docking simulations.

In this work, we performed protein-ligand docking simulations between the cryo- EM structures of inward- and outward-facing OATP1B1 and 222 ligands from an experimental dataset by Karlgren et al. [5]. This dataset designated these ligands as either inhibitors or noninhibitors of OATP1B1. We incorporated both inward- and outward-facing OATP1B1 conformations since this transporter is believed to function according to the rocker-switch alternating access mechanism, where it moves from an extracellular-facing position (i.e., outward-facing) to a cytosol-facing position (i.e., inward-facing) as it transports molecules [9]. We reasoned that docking ligands to both conformations would better sample ligand interactions occurring throughout the transport process. Protein-ligand complexes produced by the docking simulations were used as structural data to train HOLIgraph. Data representations used in HOLIgraph encompass quantitative descriptors of binding interactions (distance, angle, donor, acceptor, etc.; Table S4), expanding upon the coarse-grained protein-ligand interaction features leveraged in the earlier interaction-based models for urease inhibition [11]. HOLIgraph notably outperformed conventional ligand-based models with over 90-percent balanced accuracy. Beyond this impressive performance, HOLIgraph is built upon highly interpretable data representations that offer a wealth of insight into potential inhibition mechanisms.

## Results

HOLIgraph (Fig. 1) was designed to capture intricate inter-molecular interactions between ligands and OATP1B1 to enhance inhibition prediction accuracy. Just as ligand-based models utilize specific data representations (e.g., physicochemical lig- and properties), HOLIgraph requires specific structural data representations. In brief, these HOLIgraph representations augment traditional physicochemical ligand representations with high-dimensional descriptors of both (a) ligand position with respect to the protein topography, and (b) interactions between the ligand and protein.

**Fig. 1.**
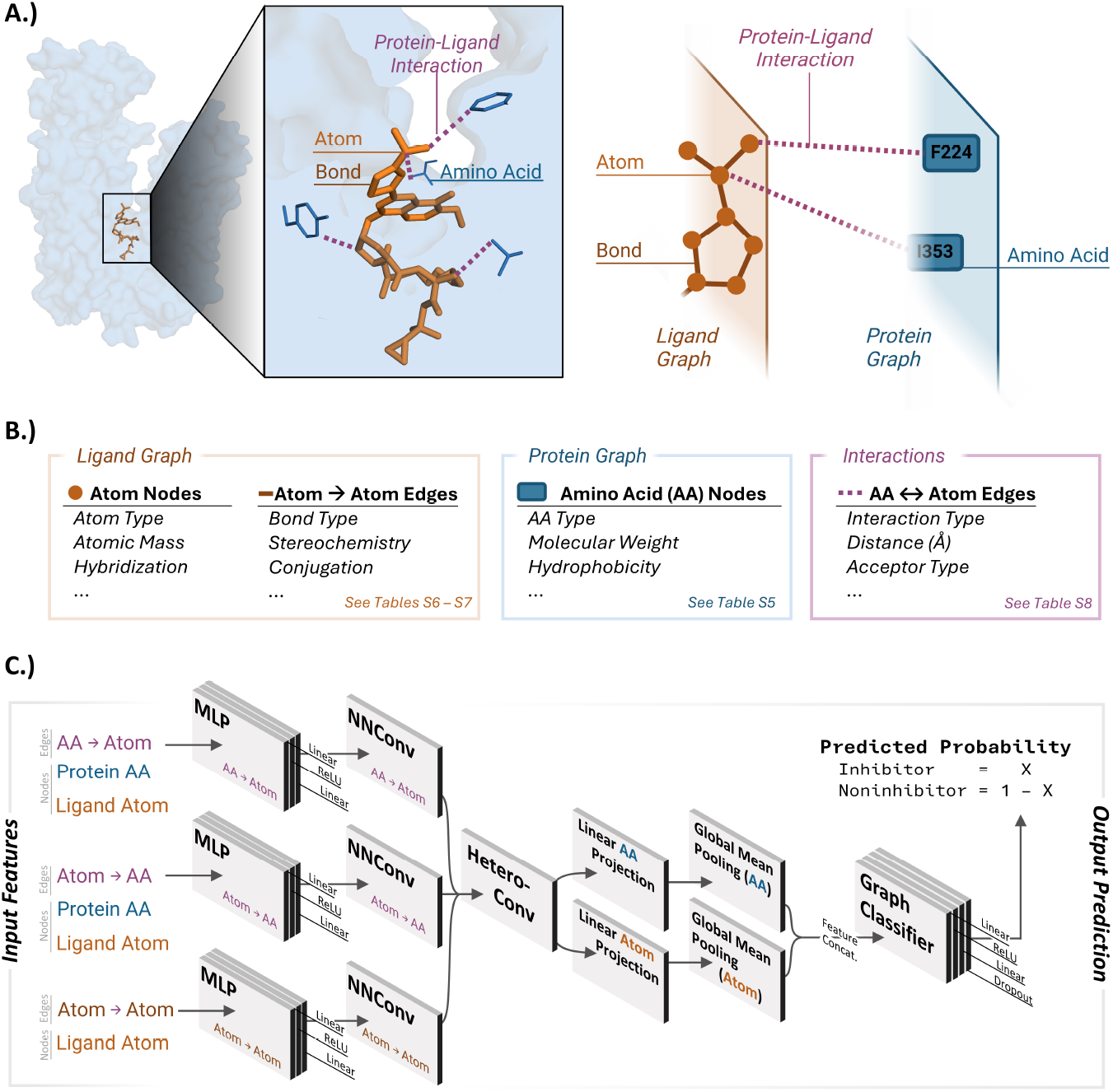
HOLIgraph encompasses a ligand graph (in orange, where the nodes are the atoms, and the edges are the intra-ligand bonds) and a protein graph (in blue, where the nodes are amino acids) connected by protein-ligand interaction edges (magenta). For example, protein-ligand interactions are shown for an OATP1B1 complex with Simeprevir, a known OATP1B1 inhibitor (A, left). A simplified HOLIgraph representation (A, right) corresponds to the interactions involving the 4-isopropylthiazole group of Simeprevir. The HOLIgraph data representation includes detailed features (B) of each atom and amino acid (AA) node, intra-ligand bond edge, and protein-ligand interaction edge. HOLIgraph features are further detailed in Tables S5–S8. The HOLIgraph architecture (C) involves processing input feature data through multilayer perceptron (MLP), neural message passing convolution (NNConv), heterogeneous graph convolution, projection, pooling, and graph classifier layers to predict the probability that a given ligand is an inhibitor or noninhibitor of OATP1B1.

### Molecular Docking Simulations Generated Novel OATP-Ligand Interaction Interfaces

Interaction interfaces between OATP1B1 and ligands were produced via protein-ligand docking simulations. Using cryo-EM structures of both the inward- and outward-facing conformers of OATP1B1 [9], we conducted thorough docking simulations to generate one thousand docked poses (i.e., protein-ligand complexes outputted by the docking simulations) for each of the 222 ligands experimentally characterized by Karlgren et al [5]. Details on the docking workflow are provided in the Supplemental Information and Fig. S2. Interaction data (e.g., interaction distances, angles, atoms involved) were extracted from each docked pose by the Protein-Ligand Interaction Profiler web tool [13]. Expanded protein-ligand interaction data is provided in the Supplemental Information (Figs. S5–S11).

### HOLIgraph Outperformed Conventional Ligand-Based Models in Predicting Ligand Activities with OATP1B1

To establish a baseline for OATP1B1 inhibitor prediction performance prior to developing HOLIgraph, we trained ligand-based classifiers on two widely used molecular representations: extended-connectivity fingerprints (ECFPs [14]) and physicochemical descriptors generated by RDKit [15] (Fig. 2A). These simple ligand-based models had modest performance, with a mean balanced accuracy of approximately 80 percent.

**Fig. 2.**
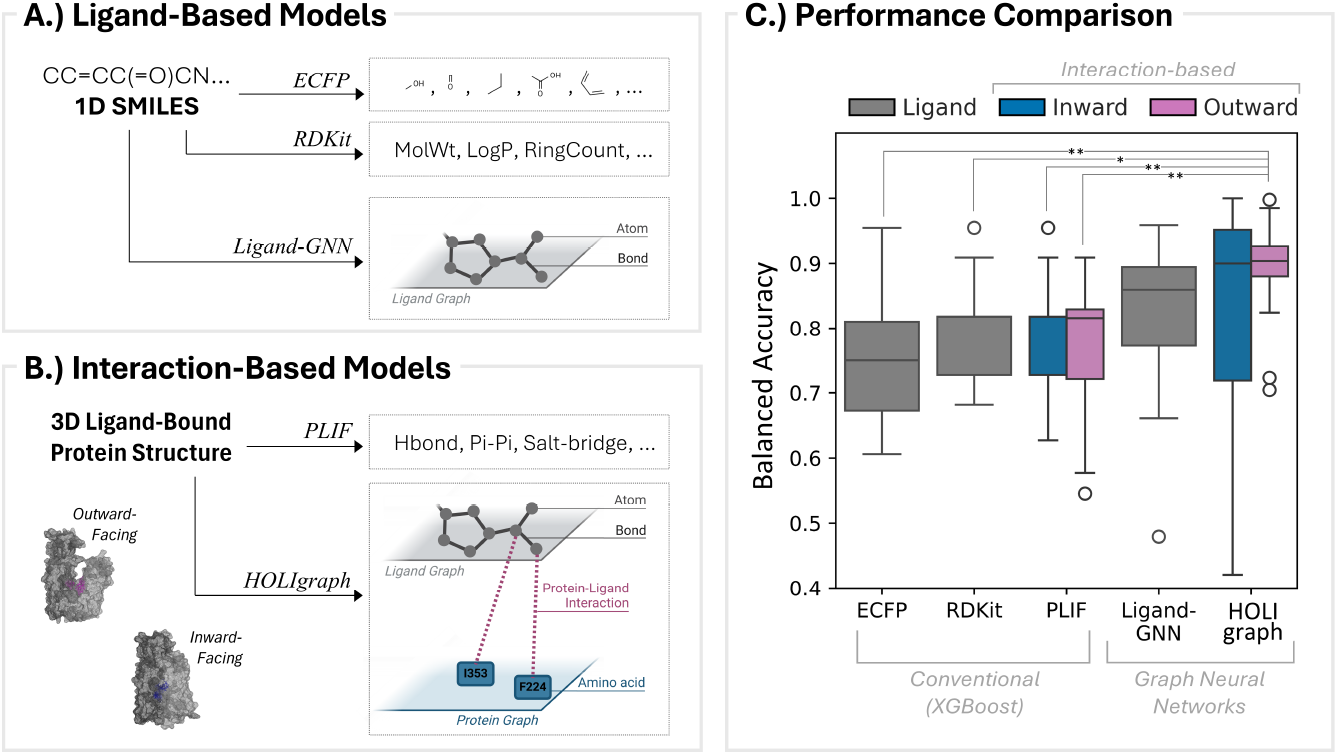
HOLIgraph outperforms ligand-based and interaction-based methods using XGBoost when trained on docked poses of outward-facing OATP1B1. (A) Feature encodings of ECFP and RDKit physicochemical descriptors generated numerical vectors and tensors for use in classification (e.g., XGBoost) and Ligand-GNN models, respectively. (B) Protein-ligand interaction fingerprints (PLIFs) obtained from docked poses were encoded into numerical vectors and tensors for use in classifier models and HOLIgraph, respectively. Feature engineering and model optimization are further detailed in the Supplemental Information. (C) Box plots displaying the distribution of scores for ligandbased (ECFP, RDKit) and interaction-based XGBoost, Ligand-GNN, and HOLIgraph (left to right). Interaction-based models for the inward- and outward-facing OATP1B1 conformers were evaluated independently (blue and pink, respectively). Mann-Whitney U p-values with Bonferroni correction indicated by asterisks (*: p*<* 0.05, **: p*<* 5e-3) show that HOLIgraph (applied to the docked poses of the outward-facing OATP1B1 conformer) improves balanced accuracy scores compared to all XGBoost models. Comprehensive performance results are reported in Tables S10–S14.

To preliminarily gauge the added value of protein-ligand interaction information, we developed simple classifiers using protein-ligand interaction fingerprints (PLIFs) (Fig. 2B, Table S4). These features captured detailed interactions between ligands and OATP1B1 in its inward- and outward-facing conformations. All models were evaluated with standard metrics including accuracy, precision, and recall (5-fold cross-validation results in Tables S10–S13). Among the classification algorithms tested, XGBoost consistently delivered the best performance for both ligand- and interaction-based models.

Building on these findings, we proceeded to develop three GNN models: a ligandbased GNN (Fig. 2A) and two interaction-based GNNs, collectively referred to as HOLIgraph, for the inward- and outward-facing conformations of OATP1B1 (Fig. 2B). HOLIgraph achieved a median balanced accuracy above 90 percent, outperforming the ligand-based GNN, which achieved 86-percent balanced accuracy. Further comparisons revealed that HOLIgraph, when using the outward-facing conformer, significantly outperformed both the ligand-based and simple interaction-based classifier models (Fig. 2C). This highlights the distinct advantage of incorporating interaction features into GNN architectures. In contrast, the ligand-based GNN did not exhibit similar improvements over conventional classifiers, underscoring the critical role of protein-ligand interaction data in enhancing predictive performance. Detailed performance metrics for the GNN models are presented in Table S14.

### Interaction data enabled comparisons of residue interactions with inhibitors versus noninhibitors

We identified critical interactions that differentiate inhibitors from noninhibitors, which showed promising agreement with recent experimental findings [9]. Fig. 3 (left) highlights the frequency of any interaction type occurring with each orthosteric site residue (Y352, F356, and F386) identified by Shan et al. [9] through cryo-EM and functional investigations. The orthosteric site (orange in Fig. 3) was defined by Shan et al. to be the set of OATP1B1 residues experimentally observed to participate in interactions with all four ligands for which they solved OATP1B1-bound cryo-EM structures. For docked poses with the outward-facing OATP1B1 conformer (Fig. 3A), inhibitors were observed to have significantly more interactions with F356 than noninhibitors (p<5E-3). Similarly, the distribution for the inward-facing OATP1B1 conformer (Fig. 3B) showed each phenylalanine residue in the orthosteric site (F356 and F386) interacted more frequently with inhibitors than noninhibitors (p< 5E-4).

**Fig. 3.**
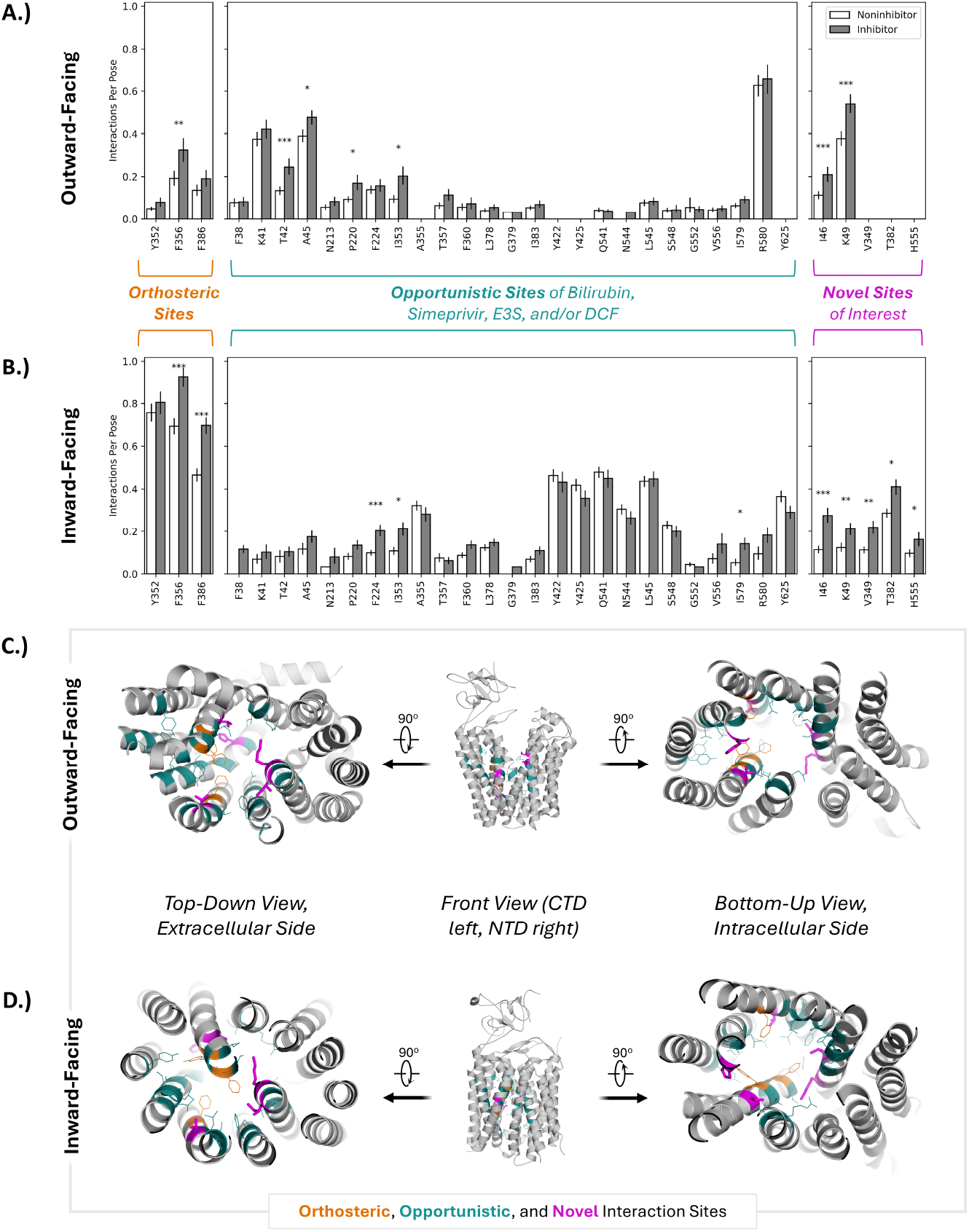
Protein-ligand interaction profiles highlight significant differences in OATP1B1 interaction sites between inhibitors and noninhibitors, including five residues not previously identified as positions of interest. Probability of any type of interaction occurring between noninhibitors (white) or inhibitors (grey) and OATP1B1 residues in the previously identified orthosteric (orange) and opportunistic (green) sites, and in novel sites of interest for the outward-facing (A) and inward-facing (B) conformers. Probabilities independently normalized for each class. Error bars indicate 95% confidence intervals. Mann-Whitney U p-values with Bonferroni correction indicated by asterisks (*: p*<* 0.05, **: p*<* 5e-3, ***: p *<* 5e-4). Orthosteric, opportunistic, and novel site residues are highlighted in the cryo-EM structures of the outward-facing (C) and inward-facing (D) conformers (PDB 8HNB and 8HND, respectively (9)). Structures are displayed in three views, from left to right: top-down (from the extracellular side), front (CTD left, NTD right), and bottom-up (from the intracellular side).

We also visualized interaction distributions for any residues in the experimentally determined opportunistic sites defined by Shan et al. to be groups of residues important for the binding of only certain ligands (i.e., much more ligand-specific than orthosteric sites) [9]. The ligands interacting with these opportunistic sites in the cryo-EM structures obtained by Shan et al. [9] were bilirubin, Simeprevir, estrone-3-sulfate (E3S), and 2’,7’-dichlorofluorescein (DCF). It is worth noting that opportunistic site residues participating in interactions indicative of ligand class (i.e., inhibitor, noninhibitor) differ greatly between the inward- and outward-facing OATP1B1 conformers. For inward-facing poses, residues F224, I353, and I579 participated in significantly more interactions with inhibitors than with noninhibitors. For outward-facing poses, residues T42, P220, and I353 were observed to have significantly more interactions with inhibitors than with noninhibitors.

Beyond interactions with orthosteric and opportunistic sites, we observed that inhibitors interacted with five additional residues considerably more than noninhibitors (Fig. 3, magenta). In particular, I46 and K49 were seen to interact more often with inhibitors than noninhibitors in both the outward- and inward-facing conformers. These findings illustrate the utility of interaction-based methods for elucidating intricate interaction patterns (Fig. S5–S10) that can inspire both predictive models and further mechanistic studies. For instance, OATP1B1 inhibitors were seen to have significantly more hydrophobic interactions with residues T42, A45, I46, K49, and P220 than noninhibitors (Fig. S5-S6). This observation aligns with the findings of Shan et al. [9], highlighting the hydrophobic packing and constriction network(s) involved in the binding, transport, and/or substrate-dependent inhibition. Moreover, comparison of interaction profiles between the inward- and outward-facing conformers yields suggestions for future mechanistic studies (e.g., functional mutagenesis studies to investigate the importance of positions I46 and K49 in OATP1B1 inhibition).

To assess agreement between cryo-EM structures [9] and docking-derived proteinligand interactions, we compared their residue interaction profiles (Fig. S11). Specifically, we examined two ligand-bound cryo-EM structures of OATP1B1: an outward-facing conformation with Simeprevir (PDB: 8HNH) and an inward-facing conformation with estrone-3-sulfate (PDB: 8HND) [9]. After tabulating each experimentally observed intermolecular interaction, we compared them to those generated from our docking workflow. Notably, the single best docking pose with Simeprevir (best in terms of interface binding energy) reproduced 70% of these experimentally observed interactions. The 30 best docking poses for estrone-3-sulfate captured all of the experimentally observed interactions, and the 30 best docking poses for Simeprevir captured all but one of the experimentally observed interactions. Thus, by expanding our analysis to include a small ensemble of docked poses, we were able to more thoroughly capture experimentally validated interactions. Moreover, this broader consideration revealed several OATP1B1 residues that have not yet been characterized but may be critical for binding. These findings support our hypothesis that ensembles of docked poses can uncover a wider array of potential interactions that may be missed by static experimental structures. However, because these conclusions currently rely on a limited number of ligands for which cryo-EM data are available, their broader applicability remains to be established.

## Discussion

Machine learning techniques have been extensively applied to predict interactions between drugs and OATPs, primarily being used in binary classifications (e.g., inhibitor/noninhibitor). Previous models for predicting OATP1B1 inhibition [5–8, 16, 17] concentrated on drug attributes and protein sequence features, with no information on protein structure or protein-ligand interactions. The exclusion of insightful, structure-based features in past models is likely due to the historical lack of experimental OATP structural information. Enabled by the exciting recent publication of several experimentally solved OATP1B1 structures (apo, holo, multiple conformers) [9], we developed the structure- and interaction-based HOLIgraph model to predict OATP inhibition. We have engineered a heterogeneous graph representation for the OATP1B1-ligand binding interface—based on intricate patterns in docking-derived protein-ligand interactions—in our novel HOLIgraph model. The improved predictive performance of HOLIgraph compared to ligand-based approaches supports our hypothesis that model augmentation with protein interaction data enables significant gains in predictive power and reveals important structural determinants of OATP1B1 inhibition.

### Interaction-Guided Hypotheses for Inhibition Mechanism

Recent experimental structures have begun to clarify the molecular mechanisms of OATP1B1, though some uncertainties remain. First, while kinetic studies have characterized competitive inhibition or suggested allosteric mechanisms for certain OATP inhibitors, exact allosteric sites have yet to be proven [1]. Thus, we allowed our ligand docking simulations to explore beyond the known major and minor binding pockets (e.g., transmembrane helices, transport channel openings), though we excluded the disordered extracellular regions (which had a mean B-factor of 82.64). We reasoned that this would also allow ligands to engage in potential allosteric interactions with OATP1B1. Despite the fact that we used a sizable simulation space during the docking, the 30 most energetically favorable binding poses for each ligand were observed to occur within the main binding region seen in the cryo-EM structures (Fig. S11). These findings provide confidence in our docking workflow and the models developed from it. Secondly, the conformational dynamics of OATP1B1 inhibition are not well understood. For instance, it is unknown whether inhibitors block transport by directly preventing conformational changes required for the alternating access mechanism (e.g., outward- to inward-facing conformational change) or by occupying a key binding site without interfering with the conformational cycle. Evidence from related solute carrier proteins suggests that inhibition can involve conformationally prohibitive mechanisms. For example, dilazep, a drug used to treat renal disorders, is known to occupy the central cavity of hENT1, thereby blocking the outward-to-inward conformational change necessary for transport [18]. Because of uncertainty in the conformational dynamics of OATP1B1 inhibition, we developed distinct models for both the outward- and inward-facing conformations. The outward-facing HOLIgraph model performed best, suggesting that key ligand interactions in the outward-facing state are influential in OATP1B1 inhibition.

Lastly, it is possible that some of the ligands explored in our study exhibit indirect OATP1B1 regulation. That is, OATP1B1 regulation not involving direct ligand interaction with OATP1B1 (e.g., transcriptional regulation). We hypothesized that the ligand-only GNN would outperform the interaction-based HOLIgraph model for ligands with such confounding interactions, as the ligand-only model excludes protein-specific information. To explore this, we assessed both Ligand-GNN and HOLIgraph performance for ligands known to interact with other transporters, enzymes, and nuclear receptors expressed in the common experimental cell line HEK293 (Fig. S4). This did not reveal any significant trends and highlights the need for comprehensive, well controlled assays. Such assays would provide datasets better suited for training mechanistically agnostic models to predict OATP1B1 inhibitors.

### Molecular docking simulations enabled extraction of extensive protein-ligand interaction information for both conformational states

These docking-derived interaction data (main findings captured in Fig. 3) leads us to several hypotheses for analyzing orthosteric sites in future studies. First, we hypothesize that Y352 is more likely involved in general ligand coordination rather than specific inhibitor binding since it shows no significant interaction difference between inhibitors and noninhibitors for both OATP1B1 conformations. Second, we hypothesize that F356 may serve a key role in stabilizing inhibitors to the outward-facing conformation of OATP1B1 since noninhibitors are less likely to interact with F356 in the outward-facing conformation compared to inhibitors. Thus, we speculate that some inhibitors bind tightly to the central cavity and prevent conformational change from the outward-facing to the inward-facing conformation.

This study also affords some insight into residues not in previously identified opportunistic or orthosteric sites. Notably, I46 is a hydrophobic residue that participated in significantly more interactions with inhibitors than with noninhibitors (Fig. 3). As I46 is known to participate in hydrophobic packing above the central OATP1B1 binding cavity for only certain substrates [9], we hypothesize nuanced rearrangements of this hydrophobic packing may engage substrate-dependent cryptic binding sites. We suggest that interactions between I46 and some inhibitors may interfere with the hydrophobic packing required for cryptic binding site availability. Similarly, our results have drawn our attention to K49, an electropositive residue that generates a charged pair with D70, thereby creating an environment amenable to substrate translocation [9]. Shan et al. have shown that the K49A mutation exhibits substrate-dependent inhibition, but that K49 does not directly interact with any of the substrates they investigated [9]. We observed that K49 forms significantly more interactions with inhibitors than with noninhibitors (in both OATP1B1 conformers), leading us to suggest a potential substrate-dependent inhibition mechanism involving disruptive interactions with K49 that prevents the electrostatic environment required for substrate translocation.

## Conclusions

From these insights, we argue that HOLIgraph and interaction-based analyses enhance our ability to understand OATP inhibition mechanisms, contributing to improved drug safety through better prediction of drug-drug interactions. However, uncertainty in OATP inhibition mechanisms add to the complexities of predictive model development. These challenges necessitate further in vitro assays and highlight the need for advanced computational methods to deconvolute nuanced data patterns. By refining our models with emerging structural and functional data, we can anticipate more accurate predictive tools to advance the drug development process.

## Methods

### Data Selection & Labeling

Several studies have been conducted over the past decade that characterize and evaluate OATP interactions with ligands [5, 6, 17, 19]. Due to notable inter-study variability among the available in-vitro OATP inhibition data [20, 21], we reasoned that sourcing data from a single set of experiments [5] would provide the most consistent framework for our study. Increased caution is warranted in data curation for inhibition modeling, as discrepancies in how inhibitors are defined in different experiments may result in misguided labeling across machine learning implementations. Experimental inhibition data (Fig. S1) was obtained from Karlgren et al. for 225 ligands assayed against OATP1B1, OATP1B3, and OATP2B1 [5]. We focused only on OATP1B1 since the other OATPs do not yet have cryo-EM structures available in both the inward- and outward-facing conformations. Of these 225 compounds, 222 were used in our workflow; we did not include 3 ligands due to conformer generation issues or obtainment errors (Table S1). As was done in the original Karlgren study [5], we labeled compounds with reported inhibition percentages greater than 50% as inhibitors (Class = 1), while we labeled those with inhibition percentages less than or equal to 50% as noninhibitors (Class = 0).

### Data Splitting for Training & Test Sets

We adopted a drug-wise splitting approach to create the holdout test sets to ensure a robust evaluation of model performance. Twenty balanced test sets were created, each containing 10% of the total dataset, with an equal representation of inhibitors and noninhibitors. The remaining 90% of the data was used for training and crossvalidation. Table S2 lists the ligands in each test set.

### Structural Modeling

Molecular docking was performed between the 222 ligands and both the outward- and inward-facing conformers of human OATP1B1 (PDB IDs 8HNB and 8HND, respectively [9]) using the Rosetta modeling suite (version 3.12) [22]. A more detailed overview of the Rosetta docking workflow can be found in the Supplemental Information. To ensure a thorough sampling of the binding space, one thousand docking simulations were performed for each protein-ligand pair, resulting in 1000 unique docked poses (protein-ligand complexes). For each docked pose, Rosetta reports the ligand-protein interface energy in Rosetta Energy Units (REU). This was used as a relative metric to rank the 1000 docked poses for each single ligand-OATP conformer pair. Poses with the lowest energy scores (i.e., highest binding affinities) were considered higher quality [23].

### Feature Engineering

Two distinct feature sets were generated: traditional ligand-only molecular descriptors (i.e., ligand features) and protein-ligand interaction features. Two types of vectorized molecular representations, extended-connectivity fingerprints (ECFP) [14] and RDKit physicochemical descriptors [15], were generated for each of the 222 docked ligands. The Protein-Ligand Interaction Profiler (PLIP) [13] was used to generate interaction profiles (XML file format) for each docked pose. PLIP produces intricate features to describe each interaction present in a docked pose, including interaction type, distance, angle, donor and acceptor atom identifiers, etc. (Table S4).

### Optimization with Simple Classification Models

Prior to training the machine learning models, preprocessing steps were applied as detailed in the Supplemental Information. Several machine learning algorithms (see Supplemental Information) were trained using the preprocessed data. Trained models were evaluated on holdout test sets. Multiple metrics were computed to assess model performance (Table S10), with area under the curve (AUC) representative of the class-weighted (i.e., balanced) accuracy score for binary classification. The number of optimal docking poses used in the HOLIgraph model was determined during classifier optimization, as further detailed in the Supplemental Information, particularly Fig. S3.

### HOLIgraph Development and Evaluation

HOLIgraph (Fig. 1) was constructed to capture the intricate interactions between the proteins and ligands. The graph contains two types of nodes: amino acid residues and ligand atoms. Edges which connect the protein and ligand graphs are referred to as heterogenous and encode interaction descriptors. HOLIgraph was trained using the AdamW optimizer with weight decay. The OneCycleLR learning rate scheduler was used for adaptive learning rates. Mixed precision training improved computational efficiency. Binary cross-entropy with logits served as the loss function. Early stopping, based on validation loss with a patience of 5 epochs, prevented overfitting. Each model trained for up to 100 epochs or until early stopping occurred. For each of the 20 data splits, a separate model was trained and evaluated on the corresponding test set. Multiple metrics were calculated: accuracy, precision, recall, F1 score, balanced accuracy, etc. This thorough evaluation provided a robust understanding of HOLIgraph’s predictive capabilities. Supplemental Information details model architecture in depth.

## List of abbreviations

*OATP:*: Organic anion transporting polypeptide
*DDI:*: Drug-drug interaction
*HOLIgraph:*: Heterogeneous OATP-Ligand Interaction Graph Neural Network
*FDA:*: Food and Drug Administration
*Cryo-EM:*: Cryogenic electron microscopy
*GNN:*: Graph neural network
*PLIP:*: Protein-Ligand Interaction Profiler
*ECFP:*: Extended-connectivity fingerprint
*PLIF:*: Protein-ligand interaction fingerprint
*XGBoost:*: Extreme gradient boosting
*hENT1:*: Human equilibrative nucleoside transporter 1
*HEK293:*: Human embryonic kidney 293
*AUC:*: Area under the curve

## Supplementary information

The accompanying Supplementary Information document is available as a separate file: Additional file 1.

## Declarations

## Ethics approval and consent to participate

Not applicable

## Consent for publication

Not applicable

## Availability of data and materials

The datasets generated and/or analyzed in this study are available in the WoldringLabMSU/HOLIgraph GitHub repository, http://github.com/WoldringLabMSU/HOLIgraph. The software required for this work is also in the WoldringLabMSU/HOLIgraph GitHub repository, available under the Creative Commons Attribution-NonCommercial-NoDerivatives 4.0 International Public License (“Public License”) at http://github.com/WoldringLabMSU/HOLIgraph.

## Authors’ contributions

B.J.O., E.M.S., and D.R.W., conceived the project; M.M., J.N.E., B.H., and D.R.W. designed research; J.N.E., M.M., T.B., N.P., A.A., and D.R.W. analyzed data and wrote the paper; all authors contributed to discussions about the project and editing the manuscript prior to submission.

## Acknowledgments

Much of the computational work in this project was performed on resources managed by the Michigan State University Institute for Cyber-Enabled Research.

