## Supplemental Information for "Predicting Inhibitors of OATP1B1 via Heterogeneous OATP-Ligand Interaction Graph Neural Network (HOLIgraph)"

### Data Selection, Labeling, & Test Sets

There is a growing recognition of the significant variability in OATP1B1 inhibition data [1, 2] which prompted us to source our data from a single study with consistent experimental conditions and labeling: Karlgren et al. (2012) [3]. Following the binary labeling scheme of the original Karlgren study, ligands with OATP1B1 inhibition percentages over 50% were deemed inhibitors (Class=1) and the remainder were deemed noninhibitors (Class=0). The distribution of inhibition measurements from Karlgren et al [3] are provided below in Fig. S1 for OATP1B1, with a dashed vertical line indicating this 50-percent inhibition cutoff. Any ligands excluded from our workflow are listed in Table S1, and all twenty of our randomly generated hold-out sets are provided in Table S2.

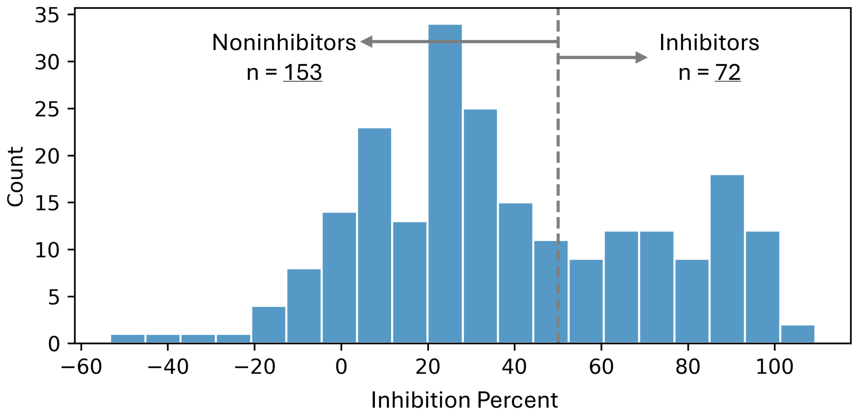


**Fig. S1** In vitro experiments resulted in the classification of 225 compounds as either inhibitors or noninhibitors of OATP1B1. The distribution of experimental inhibition percentages for all the compounds in the Karlgren et al. dataset tested against OATP1B1 [3] is shown. The 50% inhibition cutoff used to classify compounds as inhibitors is indicated by a gray dashed line. Imbalanced data prompted the use of balanced accuracy for evaluating the predictive performance of our models.

**Table S1** Ligands excluded from our dataset.

| Compound Name | Reason for Exclusion |
|---|---|
| 17-$\beta$-estradiol | Could not be obtained |
| Quinine | Same atom connectivity as quinidine (which was already obtained) in SDF file |
| Oxaliplatin | Ionic compound—conformer generation unavailable |

**Table S2**: Holdout Test Set Compositions.

| Set | Ligands |
|---|---|
| 1 | Quinidine, Chlorprothixene, Topotecan, Methotrexate, Digoxin, Ofloxacin, Flupenthixol, Allopurinol, Glycyl proline, Nefazodone, Tetracycline, Bromosulfalein, Spironolactone, Erythromycin, Indocyanine green, Paclitaxel, Repaglinide, Glycyrrhizic acid, Dipyridamole, Vinblastine, GF120918 (Elacridar), Pravastatin |
| 2 | Tranylcypromine, Fluvoxamine, Nicotine, Thioridazine, Colchicine, Enalapril, Etoposide, Eletriptan, Baicalin, Bestatin, Naringenin, Indocyanine green, Ketoconazole, Rosiglitazone, Glycochenodeoxycholate, MK-571, Candesartan, Bromosulfalein, Amprenavir, Lovastatin, Quercetin, Dipyridamole |
| 3 | Sulfaphenazole, Probenecid, Chelerythrine, Cetirizine, Terfenadine, Amantadine, Atenolol, Methoxsalen, Bestatin, Naringenin, Flupenthixol, Reserpine, Paclitaxel, Pitavastatin, Valsartan, Clarithromycin, Telmisartan, Repaglinide, Nystatin, 5-Carboxyfluorescein diacetate, Diazepam, Candesartan |

*Continued on next page*

| Set | Ligands |
|---|---|
| 4 | Erlotinib, Carbamazepine, Sildenafil, Itraconazole, Amodiaquine, Carnitine, Bufuralol, N-methylpyridinium ASP+, Triazolam, Ticlopidine, 1-methyl-4-phenyl pyridinium, Morin, Candesartan, Repaglinide, Telmisartan, Rifamycin SV, Glycyrrhizic acid, Rosuvastatin, Olmesartan, Glycodeoxycholate, Sulfasalazine, N-methylnicotinamide |
| 5 | Baicalin, Amodiaquine, Probenecid, Naringin, Midazolam, Tenofovir, Levothyroxin, Vincristine, Fluoxetine, Eletriptan, Astemizole, Nicardipine, Lopinavir, Dipyridamole, Simvastatin, Benzbromarone, Morin, Genistein, Ivermectin, Cerivastatin, Amprenavir, Pitavastatin |
| 6 | Astemizole, Erlotinib, Chloroquine, Mitoxantrone, Enalapril, Buspirone, Dofetilide, Pindolol, Diltiazem, Sotalol, Verapamil, Nifedipine, Coumestrol, Gemfibrozil, Tipranavir, Olmesartan, Cerivastatin, Rosiglitazone, Genistein, Clarithromycin, Ivermectin, Taurocholate |
| 7 | Metoprolol, Caffeine, Terfenadine, Sildenafil, Etoposide, Phenylbutazone, Methotrexate, Cimetidine, Amitriptyline, Propranolol, Theofylline, Cyclosporin, Glycocholic acid, Cerivastatin, Taurodeoxycholate, Olmesartan, Estrone-3-sulphate, Indocyanine green, Taurochenodeoxycholate, Rifampicin, Benzbromarone, Nelfinavir |
| 8 | Sanguinarine, Hoechst 33342, Triazolam, Pindolol, Tenofovir, Cimetidine, Prednisolone, Doxazosin, N-methylpyridinium ASP+, Furafylline, Methotrexate, GF120918 (Elacridar), Amprenavir, Taurochenodeoxycholate, PSC833 (Valspodar), Telmisartan, Nifedipine, Glycyrrhizic acid, Diazepam, Ketoconazole, Dipyridamole, Spironolactone |

*Continued on next page*

| Set | Ligands |
|---|---|
| 9 | Dofetilide, Isradipine, Daidzein, Cefadroxil, Paroxetine, Cholic acid, Theofylline, Tolbutamide, Prazosin, Digoxin, Verapamil, Simvastatin, Cerivastatin, Ritonavir, Ouabain, Dipyridamole, Glycochenodeoxycholate, Bromosulfalein, Indinavir, Erythromycin, Nifedipine, Spironolactone |
| 10 | Loperamide, Prazosin, Naringenin, Mephenytoin, Pindolol, Ondansetron, Sulfaphenazole, Berberine, Astemizole, Furosemide, Fentanyl, Taurodeoxycholate, Rifamycin SV, Estradiol-17-$\beta$-glucuronide, Taurolithocholate, Glycyrrhizic acid, 5-Carboxyfluorescein diacetate, Repaglinide, Olmesartan, Diazepam, Novobiocin, Fluo-3 |
| 11 | Prednisolone, Chlorzoxazone, Fexofenadine, Desipramine, Cetirizine, Procainamide, Itraconazole, Triazolam, Phenytoin, Diltiazem, Chlorprothixene, Quercetin, PSC833 (Valspodar), Diethylstilbestrol, Estrone-3-sulphate, Novobiocin, 5-Carboxyfluorescein diacetate, Rifamycin SV, Glycyrrhizic acid, Paclitaxel, Ouabain, Ezetimibe |
| 12 | Vincristine, Diltiazem, Allopurinol, Paroxetine, Glycyl proline, Testosterone, Chlorpromazine, Fexofenadine, 1-methyl-4-phenyl pyridinium, Eletriptan, Doxazosin, Lovastatin, MK-571, GF120918 (Elacridar), Amprenavir, Glycochenodeoxycholate, Taurodeoxycholate, Pitavastatin, Indinavir, Diazepam, Coumestrol, Progesterone |
| 13 | Cefamandole, Topotecan, Nitrofurantoin, Loperamide, Isoniazid, Furafylline, Erlotinib, Levothyroxin, Zidovudine, Varenicline, Captopril, Spironolactone, KO143, Nicardipine, Mifepristone, Ivermectin, PSC833 (Valspodar), Ritonavir, Tipranavir, Budesonide, Repaglinide, Genistein |

*Continued on next page*

| Set | Ligands |
| --- | --- |
| 14 | Nefazodone, Terfenadine, Vincristine, Theofylline, Celecoxib, Digoxin, N-methylpyridinium ASP+, Emtricitabine, Enalapril, 1-methyl-4-phenyl pyridinium, Ibuprofen, Pitavastatin, Saquinavir, Cholecystokinin 8, Bromosulfalein, N-methylnicotinamide, Simvastatin, Progesterone, KO143, Nifedipine, Diazepam, Benzbromarone |
| 15 | Theofylline, P-aminohippuric acid, Fluvoxamine, Topotecan, Mephenytoin, Zidovudine, Lansoprazole, Atomoxetine, Digoxin, Quinidine, Fendiline, Fluo-3, Vinblastine, Dipyridamole, Rosiglitazone, Budesonide, Cholecystokinin 8, Coumestrol, Tipranavir, Valsartan, Reserpine, Ivermectin |
| 16 | Midazolam, Celecoxib, Sanguinarine, Cefamandole, Itraconazole, Loperamide, Doxorubicin, Efavirenz, Ranolazine, Dofetilide, Chloroquine, Fluvastatin, GF120918 (Elacridar), Paclitaxel, KO143, Novobiocin, N-methylnicotinamide, Indinavir, Indometacin, Bromosulfalein, Candesartan, Glycyrrhizic acid |
| 17 | Chlorpromazine, Chelerythrine, Theofylline, Pindolol, Nicotine, Pioglitazone, Ranolazine, Buspirone, Phenylethyl isothiocyanate, Dofetilide, Piroxicam, Glycocholic acid, Budesonide, Pravastatin, Bromosulfalein, Taurocholate, Novobiocin, Atorvastatin, Taurochenodeoxycholate, Dipyridamole, Rosiglitazone, Clarithromycin |
| 18 | Penicillin G, Glipizide, Naringin, Propranolol, Glycyl proline, Sanguinarine, Theofylline, Carbamazepine, Varenicline, Isradipine, Phenobarbital, Reserpine, Mifepristone, Estradiol-17-$\beta$-glucuronide, Pravastatin, Fluvastatin, Erythromycin, Taurochenodeoxycholate, Vinblastine, Ouabain, Silymarin, Genistein |

*Continued on next page*

| Set | Ligands |
|---|---|
| 19 | Verapamil, Chlorzoxazone, Cimetidine, Tenofovir, Amitriptyline, Levothyroxin, Tolbutamide, Enalapril, Captopril, Theofylline, Amprenavir, Spironolactone, Rifamycin SV, Clarithromycin, Atorvastatin, Tipranavir, Pitavastatin, Estrone-3-sulphate, Saquinavir, Glycocholic acid, Rifampicin |
| 20 | Metoprolol, Ticlopidine, Phenylbutazone, Fentanyl, Methotrexate, Fluoxetine, Imipramine, Chloroquine, N-methylpyridinium ASP+, Acarbose, Carnitine, Novobiocin, Glibenclamide, Vinblastine, Fluvastatin, Tipranavir, Sulfasalazine, Reserpine, Genistein, Nystatin, GF120918 (Elacridar), PSC833 (Valspodar) |

### Ligand-Protein Docking Workflow

#### Ligand Structure Preparation

Canonical SMILES were compiled from PubChem [4] for subsequent conversion to three-dimensional structures (SDF format), energy minimization, and protonation (pH 7.5) using Open Babel [5]. A total of 250 conformers were generated for each ligand with the BioChemical Library conformation sampling tool [6].

#### Protein Structure Preparation

The protein sequence of human OATP1B1 (UniProtKB Q9Y6L6) was obtained from the UniProt database in FASTA format [7]. Cryo-EM structures of both the inward-facing and outward-facing OATP1B1 conformers (PDB IDs 8HND and 8HNB respectively) were obtained from the Protein Data Bank (PDB) [8]. Structures were prepared for protein-ligand docking by first removing water molecules, ions, and small molecule ligands from them manually using PyMOL [9]. The structures were then energetically minimized using the Rosetta FastRelax protocol [10] which relieves minor sidechain and backbone clashes that may be present in structures obtained from the PDB.

#### Molecular Docking Simulations

The ligand-protein docking workflow is summarized visually in Fig. S2. Python scripts were used to organize and prepare files for the Rosetta protein-ligand docking workflow [11–13]. The RosettaLigand score function was used due to its proven performance in small molecule docking compared to other Rosetta score functions and external docking algorithms [13].

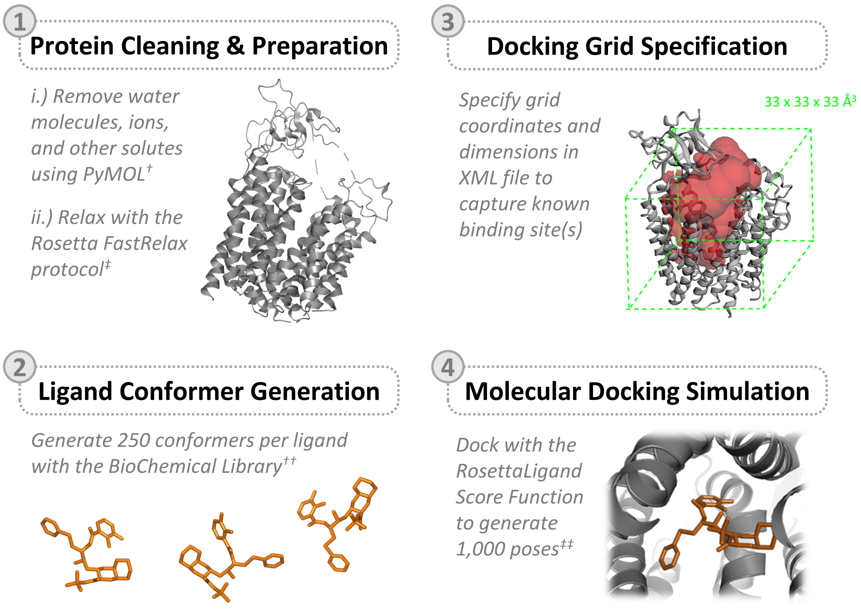


**Fig. S2** Overview of the protein-ligand docking protocol including protein structure preparation with †PyMOL [9] and ‡Rosetta FastRelax [10]; ligand conformer generation using the ††BioChemical Library [6]; docking grid specification (typical edge length 30-40 Å for OATP1B1); and docking simulations with the ‡‡RosettaLigand score function [11–13]. Figure in panel 3 created, in part, using CASTp [14].

In brief, the docking workflow involved the following critical steps: concatenating the ligand conformer files with the prepared protein structures, defining the ligand starting coordinates and the space in which ligands are allowed to move during the simulations in an XML file [13], and running the docking simulations for all ligands.

Once all the protein-ligand docking simulations were finished, the interface binding energy distributions of inhibitors and noninhibitors were analyzed. To determine if there were appreciable differences in the interface binding energy distributions of inhibitors and noninhibitors docked to the inward-facing and outward-facing conformers of OATP1B1, three statistical tests were performed (Fig S3A. shows the energy distributions for the different conformers, and Table S3 shows the statistical results).

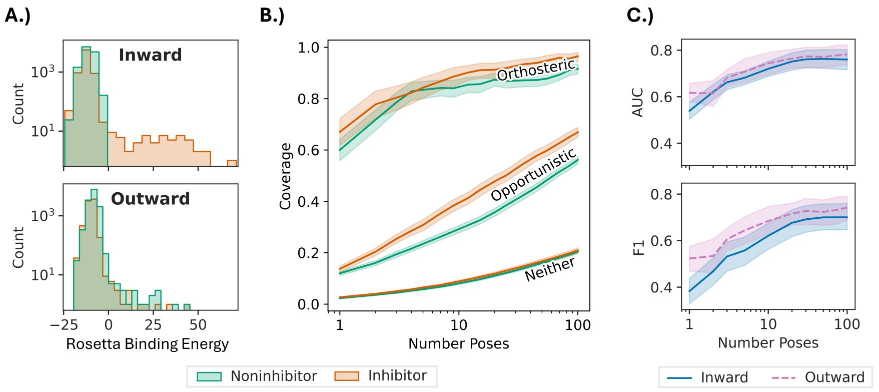


**Fig. S3** Utilizing multiple docked poses improves model performance, increases coverage of important residues, and reveals differences between the ligand classes; however, it also increases noise. (A) Distribution of Rosetta binding interface energy scores (in Rosetta energy units) of the best 100 poses for each ligand docked to the inward- and outward-facing OATP1B1 conformers (top and bottom panels for A, respectively). (B) Fraction of unique residues from the orthosteric site, opportunistic sites, and the remaining protein sequence captured in interaction descriptors as a function of number of poses. Experimentally characterized noninhibitors shown in green, and inhibitors in red. (C) Performance of the interaction-based XGBoost classification models (non-GNN) for the inward (blue, solid) and outward (pink, dashed) as a function of the number of docked poses used. Improvement in model performance indicated by AUC (top) and F1 (bottom) plateaus at approximately 30 poses. Shaded regions denote 95% confidence intervals. See Table S3 for statistical analysis.

Two methods were used to determine the statistical significance of the distribution differences: the Mann-Whitney U test [15] and the Kolmogorov-Smirnov test [16] Additionally, the practical significance of these differences was quantified by effect size (Cohen's d) [17]. These findings show that the binding energy differences between inhibitors and noninhibitors for each OATP1B1 conformer have great statistical power (i.e., very small p-values), although the magnitudes of the differences (i.e., practical significances) are fairly small. Conversely, there are large effect sizes for the differences between the energy distributions of the inward and outward OATP1B1 conformers for each compound class (i.e., inhibitors or noninhibitors), but these differences are statistically insignificant.

**Table S3** Statistical analyses corresponding to the binding energy distributions in Fig. S3.

| Groups for comparison | Mann-Whitney† | | Cohen's | Kolmogorov-Smirnov‡ | |
|---|---|---|---|---|---|
| | U-stat | p-value | d†† | KS-stat | p-value |
| Inhibitors vs. noninhibitors (inward) | 3.9E7 | 4.4E-196 | 0.27 | 0.24 | 1.0E-244 |
| Inhibitors vs. noninhibitors (outward) | 3.4E7 | 0.0 | 0.54 | 0.25 | 2.3E-265 |
| Inward vs. outward (inhibitors) | 4.7E7 | 0.0 | 0.68 | 0.48 | 0.0 |
| Inward vs. outward (noninhibitors) | 1.4E8 | 0.0 | 1.03 | 0.42 | 0.0 |

†Mann-Whitney U test is a non-parametric test to determine whether there is a significant different between the mean of two non-normal distributions. Larger U-statistic values (upper limit 8.9E9 for our data) support the null hypothesis that there is no difference between the means, whereas smaller U-statistic values (lower limit 0) supports the alternative [15].
††Cohen's d is a measure of practical effect size—that is, the difference between two means normalized to standard deviation. Cohen's d is typically interpreted as indicated small (d=0.2), medium (d=0.5), and large (d=0.8) differences between group means [17].
‡Kolmogorov-Smirnov test is a non-parametric test to compare the difference between the distributions of two groups (distribution shape, median, variability, etc). The KS-statistic ranges from zero to one, zero indicating a very small difference between distributions and one indicating a very large difference [16].

### Model Development

#### Feature Engineering

To represent OATP1B1-ligand interactions of each docked pose as protein-ligand interaction fingerprints (PLIFs), the Protein-Ligand Interaction Profiler web tool (PLIP) was employed [18]. Each docked pose was entered into PLIP, which produced an extensive interaction report. Python scripts (available at github.com/WoldringLabMSU/HOLIgraph) were used to parse interaction data from these reports into the PLIFs listed in Table S4.

**Table S4** Protein-ligand interaction fingerprint (PLIF) features.

| Feature | Description |
|---|---|
| Interaction Type | Hydrophobic, halogen/hydrogen bond, salt bridge, pi-cation, pi-stacking |
| Residue Type | One-hot encoded vector, protein residues involved in the interaction |
| Distance | Normalized distance between interacting protein residue and ligand atom |
| Angle | Normalized angle measurements (if applicable, e.g., in hydrogen bonds) |
| Donor/Acceptor Type | One-hot encoded vector, donor/acceptor roles (halogen/hydrogen bonds) |

#### HOLIgraph Architecture

The HOLIgraph architecture (Fig. 1C) is designed to model protein-drug interactions and perform graph classification tasks. It uses separate convolution layers for amino acid-to-atom, atom-to-amino acid, and atom-to-atom interactions. Edge features are first processed through multi-layer perceptrons (MLPs) to capture interaction characteristics. Convolutional layers (i.e., NNConv) then update node features based on neighboring nodes and processed edge features, propagating information across the protein-ligand interface. A HeteroConv layer aggregates outputs from these convolutions, combining information from different interaction types (i.e., intermolecular and intramolecular interactions). Linear projection layers then map atom and amino acid features to a common space. Global mean pooling is applied to create fixed-size representations for the ligand and protein. These pooled features are concatenated and passed through a classification network consisting of linear layers with ReLU activation and dropout. The network outputs a single value predicting the compound's inhibitor status. This architecture integrates heterogeneous information from protein-ligand complexes, considering both local interactions and global structure to predict inhibitory activity.

HOLIgraph, depicted in Fig. 1, consists of a protein graph and a ligand graph, connected by heterogeneous protein-ligand interaction edges. The protein graph contains amino acid nodes, each represented by 28 features (Table S5), including one-hot encoded residue type, molecular weight, aromaticity, isoelectric point, hydrophobicity, flexibility, and secondary structure propensities. These features capture the essential physicochemical properties of the protein residues involved in protein-ligand interactions.

**Table S5** Node features for amino acids.

| Feature | Type | Vector Length |
|---|---|---|
| One-Hot Encoded Residue | Binary Vector | 20 |
| Molecular Weight | Numerical | 1 |
| Aromaticity | Numerical | 1 |
| Isoelectric Point | Numerical | 1 |
| Hydrophobicity | Numerical | 1 |
| Flexibility | Numerical | 1 |
| Secondary Structure | Numerical Vector | 3 |

Ligand atom nodes are comprised of 79 atomic features (Table S6), including one-hot encoded atom type, number of heavy neighbors, formal charge, hybridization state, presence in rings, aromaticity, atomic mass, van der Waals radius, covalent radius, chirality, and number of implicit hydrogens. This set of atomic features provides a detailed representation of the ligand's chemical structure and properties.

**Table S6** Node features for ligand atoms.

| Feature | Type | Vector Length |
|---|---|---|
| One-Hot Encoded Atom Type | Binary Vector | 43 |
| Number of Heavy Neighbors | Binary Vector | 6 |
| Formal Charge | Binary Vector | 8 |
| Hybridization Type | Binary Vector | 7 |
| In a Ring | Binary | 1 |
| Aromaticity | Binary | 1 |
| Atomic Mass | Numerical | 1 |
| Van der Waals Radius | Numerical | 1 |
| Covalent Radius | Numerical | 1 |
| Chirality | Binary Vector | 4 |
| Number of Implicit Hydrogens | Binary Vector | 6 |

To represent intra-ligand bonds, 10 atom-to-atom edge features (Table S7) include bond type, conjugation, presence in rings, and stereochemistry. This graph construction process captures the local and global structural information of the protein-ligand complexes, providing a comprehensive representation for subsequent analysis.

**Table S7** Edge Features for Between Ligand Atoms.

| Feature | Description |
|---|---|
| Bond Type | One-hot encoded vector for bond types (e.g., single, double, aromatic) |
| Conjugation | Binary for whether the bond is conjugated |
| Ring Status | Binary for whether the bond is part of a ring |
| Stereochemistry | One-hot encoded vector for stereochemistry (e.g., cis, trans) |

Edges in the constructed graph represent the intermolecular interactions between amino acids and ligand atoms, as well as the intramolecular interactions among atoms within the ligand. Edges which connect heterogenous nodes encode protein-ligand interactions, represented by 332 edge PLIFs, such as interaction type, distance, angle, and other geometric properties (Table S8).

Lastly, Table S9 presents the edge formulation indices and their corresponding sizes, detailing the interactions between amino acids (labeled 'aa') and atoms, as well as the bonds between atoms.

**Table S8** Edge features for interaction graph (amino acid-ligand atom).

| Feature | Description |
|---|---|
| Interaction Type | Hydrophobic, halogen/hydrogen bond, salt bridge, pi-cation, pi-stacking |
| Residue Type | One-hot encoded vector, protein residues involved in the interaction |
| Distance | Normalized distance between interacting protein residue and ligand atom |
| Angle | Normalized angle measurements (if applicable, e.g., in hydrogen bonds) |
| Donor/Acceptor Type | One-hot encoded vector, donor/acceptor roles (halogen/hydrogen bonds) |

**Table S9** Edge formulation.

| Edge Index | Vector Length |
|---|---|
| 'aa', 'interacts with', 'atom' | 332 |
| 'atom', 'interacts with', 'aa' | 332 |
| 'atom', 'bonded to', 'atom' | 12 |

#### Data Preprocessing

One-hot encoding was performed on categorical features to convert them into a binary representation suitable for the models. Variance and multicollinearity correlation thresholding were employed to remove features with low variance (<0.001) and high correlation (>0.99), respectively, reducing the dimensionality of the feature space and mitigating the risk of overfitting. Feature normalization using StandardScaler [19] was applied to ensure that all features have zero mean and unit variance, improving the convergence and stability of the machine learning algorithms.

#### Classifier Training & Evaluation

Prior to developing HOLIgraph, we explored the use of several ML algorithms including logistic regression, support vector machines (SVM), random forests, and XGBoost. The training process involved k-fold cross-validation, where the training data was split into k subsets, and the models were trained and validated k times, using a different subset for validation each time. This approach helps to assess the models' performance and stability across different subsets of the data. The trained models were then evaluated on the holdout test sets to assess their generalization performance. Metrics such as accuracy, precision, recall, F1-score, and area under the receiver operating characteristic curve (AUC-ROC) were calculated to provide a comprehensive assessment of the model's performance in predicting OATP inhibition. Example code to generate these representations and to run the HOLIgraph model are provided on GitHub (http://github.com/WoldringLabMSU/HOLIgraph).

### Model Performance

#### Classifier Results

Detailed performance metrics for conventional classifiers using protein-ligand interaction features are provided in Tables S10 and S11 (inward and outward, respectively). Similarly, the performance of simple classification models utilizing ligand-only features ECFP [20] and RDKit physicochemical descriptors [21] are provided in Tables S12 and S13, respectively. The scores provided are the medians obtained from evaluating model performance on all 20 test sets.

**Table S10** Performance metrics for the investigated conventional classification algorithms with protein-ligand interaction features for the best 100 poses for the inward-facing OATP1B1.

| Metric | Log. Regression | | Random Forest | | SVM (RBF) | | KNN | | XGBoost | |
|---|---|---|---|---|---|---|---|---|---|---|
| | Mean | Std. | Mean | Std. | Mean | Std. | Mean | Std. | Mean | Std. |
| Accuracy | 0.50 | 0.04 | 0.52 | 0.08 | 0.43 | 0.07 | 0.39 | 0.07 | 0.58 | 0.10 |
| Precision | 0.88 | 0.09 | 0.87 | 0.12 | 0.90 | 0.09 | 0.87 | 0.11 | 0.89 | 0.10 |
| Recall | 0.44 | 0.03 | 0.48 | 0.04 | 0.33 | 0.02 | 0.28 | 0.02 | 0.54 | 0.13 |
| Balanced Accuracy | 0.57 | 0.07 | 0.58 | 0.14 | 0.55 | 0.12 | 0.51 | 0.13 | 0.62 | 0.05 |
| F1 Score | 0.58 | 0.05 | 0.62 | 0.05 | 0.49 | 0.03 | 0.43 | 0.03 | 0.67 | 0.13 |

**Table S11** Performance metrics for the investigated conventional classification algorithms with protein-ligand interaction features for the best 100 poses for the outward-facing OATP1B1.

| Metric | Log. Regression | | Random Forest | | SVM (RBF) | | KNN | | XGBoost | |
|---|---|---|---|---|---|---|---|---|---|---|
| | Mean | Std. | Mean | Std. | Mean | Std. | Mean | Std. | Mean | Std. |
| Accuracy | 0.49 | 0.02 | 0.54 | 0.06 | 0.43 | 0.04 | 0.41 | 0.05 | 0.59 | 0.10 |
| Precision | 0.87 | 0.09 | 0.86 | 0.11 | 0.88 | 0.11 | 0.85 | 0.11 | 0.87 | 0.10 |
| Recall | 0.38 | 0.08 | 0.47 | 0.05 | 0.29 | 0.07 | 0.27 | 0.06 | 0.53 | 0.13 |
| Balanced Accuracy | 0.60 | 0.03 | 0.62 | 0.12 | 0.59 | 0.05 | 0.56 | 0.04 | 0.65 | 0.08 |
| F1 Score | 0.53 | 0.09 | 0.61 | 0.06 | 0.43 | 0.10 | 0.41 | 0.08 | 0.66 | 0.14 |

**Table S12** Performance metrics for the investigated conventional classification algorithms with Extended Connectivity Fingerprints (ECFPs).

| Metric | Logistic Regression | | Random Forest | | KNN | | XGBoost | |
|---|---|---|---|---|---|---|---|---|
| | Mean | Std. | Mean | Std. | Mean | Std. | Mean | Std. |
| Accuracy | 0.74 | 0.05 | 0.74 | 0.05 | 0.61 | 0.07 | 0.75 | 0.09 |
| Precision | 0.81 | 0.18 | 0.83 | 0.16 | 0.40 | 0.55 | 0.78 | 0.15 |
| Recall | 0.51 | 0.04 | 0.47 | 0.04 | 0.05 | 0.08 | 0.59 | 0.15 |
| Balanced Accuracy | 0.71 | 0.04 | 0.70 | 0.03 0.53 | 0.04 | 0.74 | 0.10 | |
| F1 Score | 0.62 0.06 | 0.60 | 0.05 | 0.10 | 0.14 | 0.66 | 0.11 | |

**Table S13** Performance metrics for the investigated conventional classification algorithms with RDKit physicochemical ligand descriptors.

| Metric | Logistic Regression | | Random Forest | | KNN | | XGBoost | |
|---|---|---|---|---|---|---|---|---|
| | Mean | Std. | Mean | Std. | Mean | Std. | Mean | Std. |
| Accuracy | 0.78 | 0.07 | 0.81 | 0.06 | 0.70 | 0.05 | 0.79 | 0.07 |
| Precision | 0.78 | 0.09 | 0.81 | 0.12 | 0.73 | 0.16 | 0.81 | 0.07 |
| Recall | 0.64 | 0.15 | 0.69 | 0.10 | 0.39 | 0.08 | 0.64 | 0.12 |
| Balanced Accuracy | 0.76 | 0.07 | 0.79 | 0.06 | 0.65 | 0.04 | 0.77 | 0.06 |
| F1 Score | 0.69 | 0.08 | 0.73 | 0.08 | 0.50 | 0.07 | 0.71 | 0.08 |

### HOLIgraph Results

Detailed performance metrics for HOLIgraph (for each OATP1B1 conformer) and Ligand-GNN are provided in Table S14. The scores provided are the medians obtained from evaluating model performance on all 20 test sets.

### Considerations for Model Evaluation

PLIF-based models like HOLIgraph rely on the assumption that OATP inhibition occurs via a direct OATP-ligand interaction (e.g., allosteric or competitive inhibition).

**Table S14** Median performance metrics for test set predictions of Ligand-GNN and HOLIgraph models.

| Metric | Ligand-GNN | HOLIgraph (Inward) | HOLIgraph (Outward) |
|---|---|---|---|
| Accuracy | 0.705 | 0.813 | 0.782 |
| Precision | 0.785 | 0.782 | 0.802 |
| Recall | 0.705 | 0.699 | 0.789 |
| AUC | 0.860 | 0.900 | 0.904 |
| F1 Score | 0.691 | 0.725 | 0.781 |

However, it is likely that not all of the inhibitors explored in our study exhibit such mechanisms. Instead, we speculate that interactions with other transporters, metabolic enzymes, and nuclear receptors may play indirect roles in the perceived OATP1B1 inhibition, or lack thereof.

We searched the KEGG Database [22–24], supplemented with several publications [25–29], to determine which ligands may participate in indirect OATP1B1 regulation. We considered interactions with four categories of proteins known to be expressed by HEK293 cells, or kidney cells generally: (i) uptake proteins OCT2, OAT1, and OAT3; (ii) efflux proteins URAT1, MATE2, P-gp, MRP1, and MRP4; (iii) cytochrome P450 3A4 (CYP3A4); and (iv) nuclear receptors implicated in transport and metabolism regulation (liver X receptor, LXR; farnesoid X receptor, FXR; pregnane X receptor, PXR) [25–28]. We found that nearly 100 ligands in our dataset have been observed to interact with one or more of the aforementioned proteins. For ligands found to have potentially confounding interactions, we compared the performance of HOLIgraph (outward-facing) and Ligand-GNN (Fig. S4), though no significant correlations were discovered.

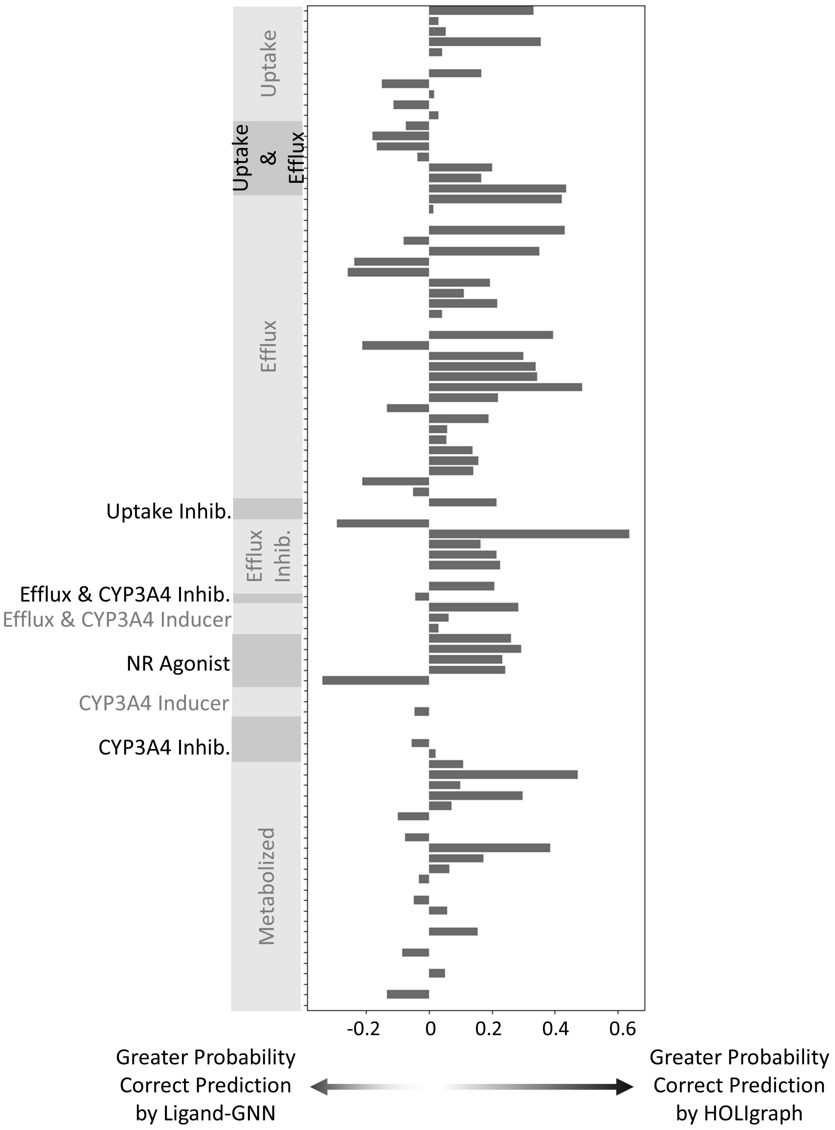


**Fig. S4** Difference in probability of correct prediction by HOLIgraph vs Ligand-GNN for ligands with potentially confounding interactions. Data shown only for ligands that appeared in three or more test/validation sets. Annotations indicate ligands with known activity that may be implicated in alternative routes of OATP1B1 modulation. This includes "Uptake" by HEK293 transporters other than OATP1B1 (OCT2, OAT1, OAT3); "Efflux" by HEK293 transporters (URAT1, MATE2, P-gp, MRP1, MRP4); CYP3A4 metabolism, or direct inhibition or induction of these processes. Also annotated are known agonists of HEK293 nuclear receptors (NRs) [25–29].

### Expanded Protein-Ligand Interaction Analyses

#### Key Residues Involved in Interactions with Inhibitors vs. Noninhibitors

Herein, we provide a breakdown of these interaction distributions by interaction type to further elucidate residue involvement in potential inhibition mechanisms. Hydrophobic interaction frequency is shown in Fig. S5.

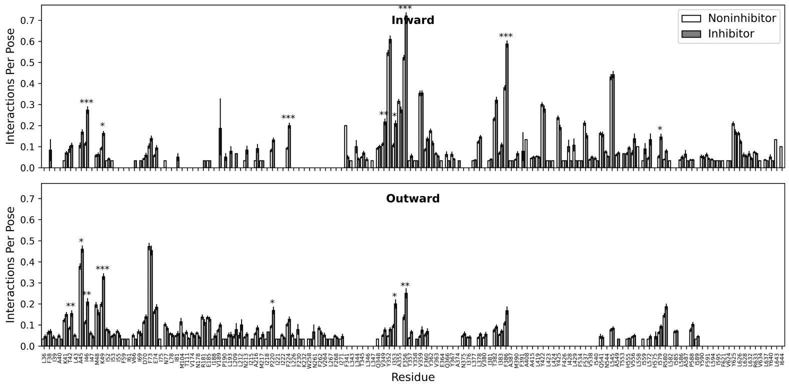


**Fig. S5** Hydrophobic interaction distribution by residue for best 30 poses of each ligand-OATP1B1 conformer pair (inward, top; outward, bottom). Noninhibitor data in white (unfilled bars); inhibitor data in grey (filled bars). Error bars indicate 95% confidence interval. Mann-Whitney U p-values with Bonferroni correction indicated by asterisks (*: $p < 0.05$, **: $p <$ 5e-3, ***: $p <$ 5e-4).

In the inward-facing conformation, eight residues were seen to interact more often with inhibitors compared to noninhibitors: I46, K49, F224, V349, I353, F356, F386, and I579 (Fig. S6, left). Similarly, inhibitors were observed to have significantly more hydrophobic interactions with seven residues in the outward-facing conformer: T42, A45, I46, K49, P220, I353, and F356 (Fig. S6, right). This supports the observation of

Shan et al [30]—who published the cryo-EM structure of OATP1B1—that hydrophobic packing plays an important role in the inhibition of OATP1B1. This motivates subsequent mutagenesis studies to investigate any role of these residues in transporter function. The statistical significance of our findings for hydrophobic interactions are based on criteria of Mann-Whitney U p-values less than 0.05 after Bonferroni correction.

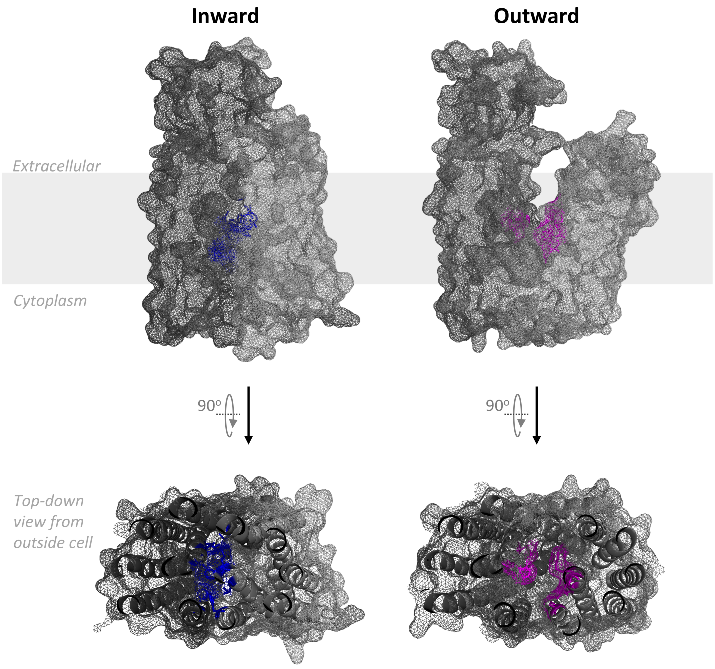


**Fig. S6** Residues observed to have significantly more hydrophobic interactions with inhibitory ligands compared to noninhibitors. Eight residues of the inward-facing conformer (left, blue) were notably more likely to participate in hydrophobic interactions with inhibitors than with noninhibitors. The outward-facing conformer (right, magenta) had seven residues displaying a greater probability of hydrophobic interactions with inhibitors. Residues I46, K49, I353, and F356 were all observed to have this increased propensity in both the inward and outward conformers.

We explored these trends for other interaction types (hydrogen/halogen bonding, pi-stacking, pi-cation interactions; Fig. S7—S10, respectively), though no rigorous statistical differences were seen for these interaction types.

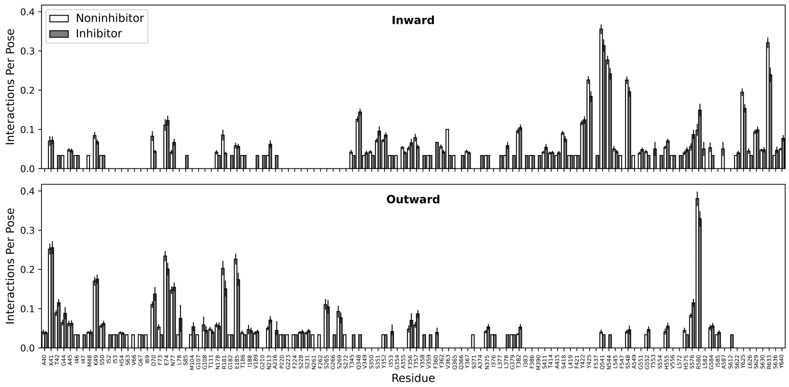


**Fig. S7** Hydrogen bond distribution by residue for the best 30 poses of each ligand-OATP1B1 conformer pair (inward, top; outward, bottom). Noninhibitor data in white (unfilled bars); inhibitor data in grey (filled bars). Error bars indicate 95% confidence interval. Mann-Whitney U p-values with Bonferroni correction found no significant differences between classes.

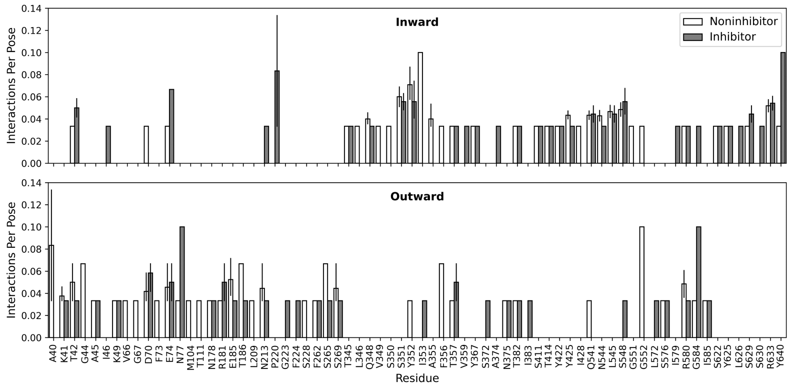


**Fig. S8** Halogen bond distribution by residue for the best 30 poses of each ligand-OATP1B1 conformer pair (inward, top; outward, bottom). Noninhibitor data in white (unfilled bars); inhibitor data in grey (filled bars). Error bars indicate 95% confidence interval. Mann-Whitney U p-values with Bonferroni correction found no significant differences between classes.

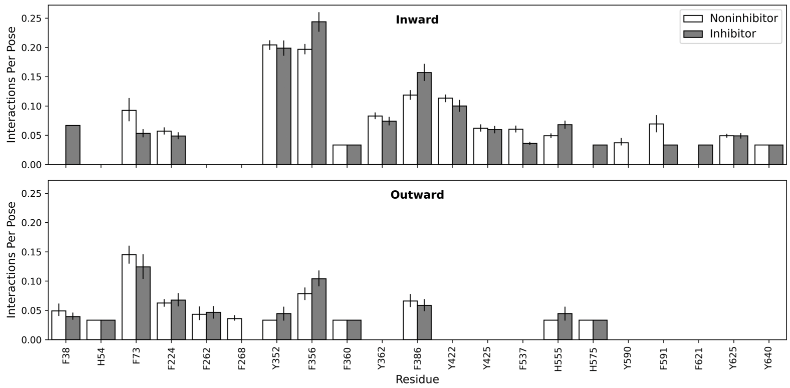


**Fig. S9** Pi-stacking interaction distribution by residue for the best 30 poses of each ligand-OATP1B1 conformer pair (inward, top; outward, bottom). Noninhibitor data in white (unfilled bars); inhibitor data in grey (filled bars). Error bars indicate 95% confidence interval. Mann-Whitney U p-values with Bonferroni correction found no significant differences between classes.

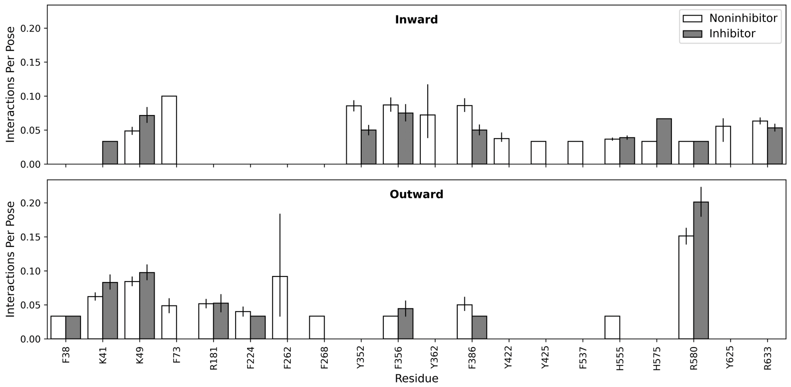


**Fig. S10** Pi-cation interaction distribution by residue for the best 30 poses of each ligand-OATP1B1 conformer pair (inward, top; outward, bottom). Noninhibitor data in white (unfilled bars); inhibitor data in grey (filled bars). Error bars indicate 95% confidence interval. Mann-Whitney U p-values with Bonferroni correction found no significant differences between classes.

#### Comparison of Docking Poses with Cryo-EM Structures

We applied our PLIF extraction workflow to the E3S-bound cryo-EM structure (PDB ID 8HND) [30] for comparison with our docking-derived PLIFs for E3S. As the experimental E3S-bound structure is inward-facing, we selected data for only our inward-facing E3S docking poses. Fig. S11A provides the cryo-EM PLIF distribution by residue in magenta, overlayed with the PLIF distributions for the single most stable docked pose (blue) and for the 30 most stable poses (orange). As expected, the dataset containing 30 poses displayed a much broader distribution than either the single cryo-EM [30] or docking structure. Since our dataset contained no other ligands with experimentally solved OATP1B1-bound structures, we repeated our docking-to-PLIF workflow for Simeprevir, as a structure was available for the outward-facing OATP1B1 bound to Simeprevir [30]. Results are provided in Fig. S11B, with findings aligned to those aforementioned for E3S.

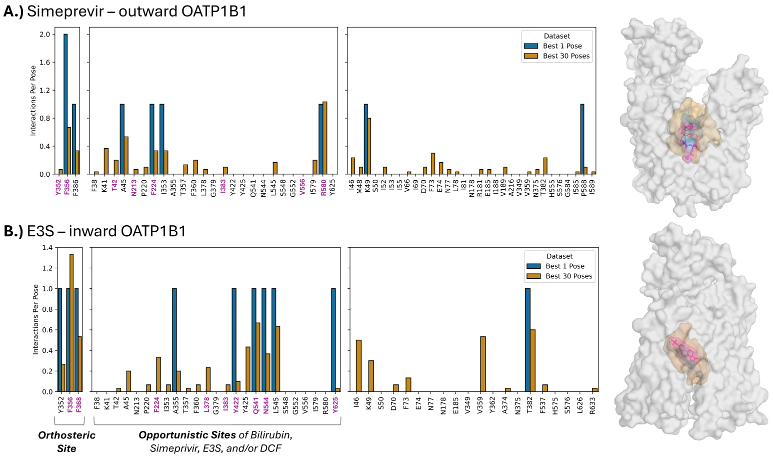


**Fig. S11** Cryo-EM structures of OATP1B1 in complex with Simeprevir and estrone-3-sulfate (27) have many of the same residue interactions as the best docking poses. Comparison of cryo-EM and docking-derived residue interactions for (A) Simeprevir bound to the outward-facing conformer and (B) estrone-3-sulfate bound to the inward-facing conformer. The number of interactions per pose was calculated by dividing the number of interactions with a given residue by the number of poses analyzed (e.g., one or thirty). The residue numbers colored in magenta indicate residues that formed interactions with either Simeprevir (A) or estrone-3-sulfate (B) in their respective cryo-EM structures. On the right of both panel A and B, cryo-EM structures by Shan et al. [30] are shown with the ligand in magenta, overlayed with a surface map of the interacting residues in the best docked pose (blue) and the thirty best poses (orange).

Using an ensemble of multiple docked poses per ligand provided a broad view of transporter interactions, both conformationally and energetically. We posit that our enhanced spatial sampling technique (i.e., using an ensemble of 30 docked poses for each ligand to train HOLIgraph) may be more representative of interactions with transporters relative to the static representations offered by single docked poses or cryo-EM structures. Comparisons between our docking-derived interaction data to those of experimentally determined, ligand-bound OATP1B1 structures [30] confirmed that our synthetic docked structures not only agreed with the experimental ligand-bound structures but also revealed a broader range of interactions. Fig. S11 presents the distribution of residues involved in protein-ligand interactions for the inward and outward cryo-EM structures bound to estrone-3-sulfate and Simeprevir, respectively, in magenta. For comparison, these distributions are also displayed for the docked poses with Simeprevir and estrone-3-sulfate, considering only the best pose (blue) and the best 30 poses (orange) for each ligand. The sampling space expansion certainly increases noise (by including interacting residues from unlikely binding poses, panel B of fig. S3), however, our simple classifier models allowed us to make great progress in striking a balance between binding site coverage and signal-to-noise ratio.

#### Multi-Pose Sampling Expands Binding Site Coverage at Expense of Signal.

Rosetta docking simulations produced 1,000 docked poses (protein-ligand complexes) of varying quality for each ligand. The Rosetta binding interface energy score was used as a quality metric, with lower energies indicating more stable poses [11]. Panel A of Fig. S3 illustrates energy distributions of the 100 most stable poses for both inhibitors and noninhibitors docked to the inward and outward conformations of OATP1B1 (upper and lower panels, respectively). Interestingly, noninhibitors docked to the inward conformer are generally more stable than inhibitors. Conversely, noninhibitors docked to the outward conformer are slightly less stable than inhibitors, although to a lesser extent (statistical support in Table S3). This may lend some support to the hypothesis that inhibitors can bind more tightly to the initial outward conformation to impair conformational change and transport, whereas noninhibitors (which include transported compounds) enable the outward-to-inward conformational change [30].

We did not use all 1000 docked poses for each protein-ligand pair to train HOLIgraph, since we hypothesized that much of the docking data contained noise. Instead, we investigated the number of poses needed to achieve optimal performance. To achieve optimal HOLIgraph performance, as indicated in Fig. 2, we applied simple machine learning engineering techniques to maximize the signal obtained from our computational modeling, effectively separating meaningful patterns from noise. This involved using both inward- and outward-facing OATP1B1 conformers and varying the number of docked poses used per protein-ligand pair. The number of docked poses used in this optimization analysis ranged from solely the best pose to the best 100 poses.

The residue-interaction distributions of our docking-derived interaction data were evaluated for consistency with experimentally identified interaction sites. Our analyses focused on three groups of OATP1B1 residues: orthosteric site residues, opportunistic sites, and residues not in the orthosteric or opportunistic sites. Specifically, we considered a broadly encompassing set of residues present in the opportunistic sites of Simeprevir, estrone-3-sulfate, bilirubin, and 2',7'-dichlorofluorescein (DCF), as done by Shan et al [30]. As expected, increasing the number of docked poses from 1 to 100 resulted in a greater diversity of OATP1B1 residues captured in the binding interactions of each ligand. Panel B of Fig. S3 demonstrates how increasing the number of docked poses increases the coverage of the orthosteric, opportunistic, and other OATP1B1 residues.

While expanding the pose sampling space may have increased the binding site coverage in the interaction profiles, it did not necessarily result in improved model performance. Prior to GNN implementation, we assessed the performance of a simpler XGBoost classification model as a function of number of poses per ligand. The performance of this simpler model (as measured by F1 score and AUC) was observed to peak when approximately 30 poses per ligand were included in the dataset for both the inward and outward conformers (panel C of Fig. S3). Thus, we used the best 30 poses for each ligand when training HOLIgraph.

#### Considerations for the Direct Inhibition Assumption

HOLIgraph relies on the assumption that inhibitors act via a direct inhibition mechanism (rather than an indirect mechanism such as transcriptional regulation), as our feature representations encompass interactions specific to the OATP1B1 transport channel. Competitive inhibition was assumed in the study by Karlgren et al. that provided the experimental labels for our model [3], as is customary in OATP1B1 inhibition assays [31, 32]. The validity of this assumption—as applied to both HOLIgraph and experimental data—has been called into question by emerging evidence of alternative mechanisms by which certain ligands may inhibit OATP function [25, 29, 33, 34].

Convolution arises from the possible involvement of metabolism, efflux, or non-OATP1B1 uptake of investigated ligands. Indirect inhibition of OATP1B1 may occur as transcriptional modulation via the farnesoid X receptor (FXR), hepatocyte nuclear factor (HNF) 4$\alpha$ and 1$\alpha$ , and liver X receptor (LXR) $\alpha$ [29]. In part, this has motivated the standard use of the human embryonic kidney (HEK) 293 cell line for in vitro OATP inhibition assays due to their low basal expression levels of many transporter proteins [25]. However, this does not consider the effect of ligand uptake on expression modulation of transporters and/or metabolic enzymes. For example, in developing kidney cells, HNF1$\alpha$ and HNF4$\alpha$ are suspected to regulate expression of several organic anion transporters (OATs) [35–37], which are known to facilitate uptake of multiple ligands in our dataset [29].

Further confounding processes include ligand-induced post-translational and/or degradative regulation. For instance, the protein kinase C activator phorbol 12-myristate 13-acetate (PMA), a compound widely studied in cancer and hematology research [38, 39], results in the phosphorylation and subsequent deactivation of OATP1B3 [40]. The extent to which any of these processes may impact in vitro OATP inhibition assays is unclear [1]. Timescales of transcriptional regulation in mammalian cells range from tens of minutes to several hours [41], while post-translational events occur in seconds. Thus, some indirect regulatory processes are theoretically plausible within the suggested 30-minute experimental pre-incubation period.

For ligands with such indirect means of OATP inhibition, we hypothesize that ligand-based models (e.g., ECFP [20], RDKit [21], and the Ligand-GNN model that we developed) may be advantageous for predicting inhibitory activity. This is because ligand-centric models are not constrained by OATP-specific information. However, for ligands whose primary mode of inhibition is direct (which is generally assumed with OATPs and related transporters), ligand-centric approaches likely cannot fully capture the intricate protein-ligand interaction features that we hypothesized to be important for determining inhibition.
